# Orientations of Mistaken Point fronds indicate morphology impacted ability to survive turbulence

**DOI:** 10.1101/2021.09.10.459851

**Authors:** Philip B. Vixseboxse, Charlotte G. Kenchington, Frances S. Dunn, Emily G. Mitchell

## Abstract

The Ediacaran organisms of the Mistaken Point E surface have provided crucial insight into early animal communities, including how they reproduced, the importance of Ediacaran height and what the most important factors were to their community dynamics. Here, we use this iconic community to investigate how morphological variation between eight taxa affected their ability to withstand different flow conditions. For each of *Beothukis, Bradgatia, Charniodiscus procerus, Charniodiscus spinosus, Plumeropriscum, Primocandelabrum* and *Fractofusus* we measured the orientation and length of their stems (if present) and their fronds. We statistically tested each taxon’s stem and frond orientation distributions to see whether they displayed a uniform or multimodal distribution. Where multimodal distributions were identified, the stem/frond length of each cohort was tested to identify if there were differences in size between different orientation groups. We find that *Bradgatia* and *Thectardis* show a bimodal felling direction, and infer that they were felled by the turbulent head of the felling flow. In contrast, the frondose rangeomorphs including *Beothukis, Plumeropriscum, Primocandelabrum*, and the arboreomorphs were felled in a single direction, indicating that they were upright in the water column, and were likely felled by the laminar tail of the felling flow. These differences in directionality suggests that an elongate habit, and particularly possession of a stem, lent greater resilience to frondose taxa against turbulent flows, suggesting that such taxa would have had improved survivability in conditions with higher background turbulence than taxa like *Bradgatia* and *Thectardis*, which lacked a stem and which had a higher centre of mass, which may have fared better in quieter water conditions.

## INTRODUCTION

The Ediacaran macrobiota is a probably polyphyletic assemblage of organisms which appear in the fossil record ∼575 million years ago and contain some of the oldest animals in the fossil record (Xiao and Laflamme 2009; Budd and Jensen 2017; Bobrovskiy et al. 2018; Cuthill and Han 2018; Dunn et al. 2018, 2021; Wood et al. 2019). The morphologies of Ediacaran organisms from Newfoundland and the UK have few clear points of homology with living animal lineages or Phanerozoic fossil groups, which has historically limited our understanding of their phylogenetic affinities and hampers our understanding of the functional ecology of these organisms (Liu et al. 2015).

The Ediacaran communities of Eastern Newfoundland are dominated by the perhaps most distinct members of the Ediacaran macrobiota - the sessile, frondose rangeomorphs (Narbonne and Gehling 2003; Narbonne 2005). Rangeomorphs are characterised by a “fractal” branching architecture (Narbonne 2004; Cuthill and Morris 2014), and which increasing data supports as a clade of stem-group eumetazoans (Cuthill and Han 2018; Dunn et al. 2021). Rangeomorphs numerically dominate these late-Ediacaran sea floors, but they lived alongside a number of different groups the most abundant of which are the arboreomorphs (Clapham et al. 2003; Xiao and Laflamme 2009). These are similarly frondose, but unlike the rangeomorphs which can possess many orders of hierarchical branching, Newfoundland arboreomorphs possess only two (Laflamme et al. 2004; Laflamme and Narbonne 2008; Laflamme et al. 2018). Non-frondose fossils are also present, though rare, in these fossil deposits and the most well-known is *Thectardis* a conical to triangular organism sometimes interpreted as a sponge (Clapham et al. 2004; Sperling et al. 2011).

Of these groups, rangeomorphs are not only the most diverse but display the greatest anatomical variation (Shen et al. 2008; Xiao and Laflamme 2009). Some rangeomorphs are preserved as single fronds (e.g. *Charnia*), but others were bushy (e.g. *Bradgatia*), spindle-shaped (e.g. *Fractofusus*) or arborescent (e.g. *Primocandelabrum*) (Gehling and Narbonne 2007; Bamforth et al. 2008; Flude and Narbonne 2008; Bamforth and Narbonne 2009; Dunn et al. 2019). Rangeomorph branches differentiated directly from one another or from a central stalk (Dunn et al. 2019) and some rangeomorphs additionally exhibited a naked stem which elevated the frond into the water column upright taxa (Laflamme et al. 2012) increasing the dispersal range of offspring (Mitchell and Kenchington 2018). Most rangeomorphs possessed a spheroidal-discoidal holdfast which anchored them within the sediment (Laflamme et al. 2004), attaching the organism to the substrate and from which the stem or frond derived. Previous functional studies have demonstrated that the high surface area of the repeatedly branched frond maximised nutrient or gas exchange (Laflamme et al. 2009; Sperling et al. 2011; Liu et al. 2015). The phylogenetic relationship between frondose rangeomorphs and the coeval arboreomorphs is currently unclear (Dececchi et al. 2017; Cuthill and Han 2018); some have argued that arboreomorphs are members of the Rangeomorpha (Brasier and Antcliffe 2009), but clear anatomical differences between at least some arboreomorphs and rangeomorphs mean that this view is not universally held, with others suggesting that overtly similar gross morphologies may have arisen through convergence (Laflamme et al. 2018). Indeed, in the modern a sessile, frondose bodyplan is found in myriad different groups, such as ferns, corals and cnidarians, and has been acquired through different developmental processes, demostrating that such a bodyplan can be the produce of similar ecologies or function and is not necessarily indicative of close phylogenetic relationship.

Stems were originally thought to facilitate height-driven tiering in Avalonian communities, allowing taller fronds to reach higher-velocity conditions (Ghisalberti et al. 2014), but more recent work has suggested that not all communities were tiered and that increased height may have additionally functioned in offspring dispersal (Mitchell and Kenchington 2018).

Thickening of the stem close to the holdfast – optimisation of the stem as a cantilever beam – is observed in cnidarians (Koehl 1977*a*, *b*), and crinoids (Baumiller and Ausich 1996), where it permits orientation of the crown with the aboral surface facing the flow, initiating aboral inflow and recirculation (Dynowski et al. 2016). Rangeomorphs have been documented as showing a basal thickening of the stem and so may have functioned in the same way (Kenchington and Wilby 2017). By examining the different functional ecology of stemmed and non-stemmed organisms, we can investigate what the advantages of stems were in Ediacaran organisms.

These fossils are found preserved within turbiditic sequences, under thin layers of ash which blanketed large swathes of sea floor and smothered thousands of macro-organisms in a single event bed (Wood et al. 2003). Communities are exceptionally preserved and provide a near-census record of the benthic communities (Wood et al. 2003). This in-situ preservation, combined with the sessile habit of the organisms, means that detailed spatial ecological analyses can be used to investigate reproductive strategies (Mitchell et al. 2015), taxonomy (Mitchell et al. 2018), community interactions (Mitchell and Butterfield 2018) and evolutionary drivers (Mitchell et al. 2019, 2020), and in this study supplement functional ecology analyses of the organisms.

Here, we use statistical analyses of the orientations of 8 taxa from the E surface, Mistaken Point, Newfoundland: *Beothukis, Bradgatia, Charniodiscus procerus, Charniodiscus spinosus, Plumeropriscum, Primocandelabrum, Thectardis*, and *Fractofusus*. We determine the extent to which orientation distributions of populations of complete specimens, stems and fronds are randomly, normally and/or uniformly distributed, and how many sub-groups within each population exist. Where taxa exhibit multi-modal orientation distributions, we use random labelling spatial analyses to determine whether there are any spatial patterns to taxa orientations. These analyses enable us to investigate how morphological features, such as stems and number of folia influenced the stability of these organisms in the ancient oceans and their ability to withstand burial events of differing magnitudes.

## GEOLOGICAL SETTING

The Avalon Assemblage records the evolution of deep marine metazoan communities from the ∼574 Ma Drook Formation (Matthews et al. 2021), to the late Ediacaran Bradgate Formation (556.6±6.4 Ma, (Noble et al. 2015)). One of the three assemblages originally proposed by Waggoner 2003, it traces the marine margin of the Avalonian Terrane through the British Isles and Newfoundland. In both regions, sedimentation was dominated by turbidite deposition (Wood et al. 2003; Noble et al. 2015). Throughout the Newfoundland succession, there is a transition in tectonic setting and depositional character, from the basin plain setting of the lower Conception Group to the shallowing-upwards slope setting of the upper Conception and St John’s groups, with a concomitant increase in depositional energy and rate of deposition and a basinwards progradation of the locus of sedimentation (Wood et al. 2003; Matthews et al. 2021).

The Mistaken Point Formation is dominated by thick-bedded, mud-rich and ashy turbidites, punctuated by tuffaceous horizons (Wood et al. 2003; Ichaso et al. 2007; Matthews et al. 2021). The bed over the E surface has a thin, coarse crystal tuff, a lower graded portion and an upper portion that consists of alternating dark-light bands (above the chlorite-carbonate band; (Fig S4; Matthews et al. 2021). The exact mode of emplacement of the tuffaceous horizons was long thought to be primarily from water-lain ashfall events (where ashy material enters the basin, and gradually settles out through the water column). However, recent work suggests that at least some of these horizons were instead the product of ashy turbidites, and that they contain variable proportions of volcaniclastic (eruptive and/or unlithified reworked) and epiclastic (lithified and reworked) material (Noble et al. 2015; Kenchington et al. 2018; Matthews et al. 2021).

The mode of emplacement has direct implications for understanding the process that felled the fronds within the palaeocommunities, and therefore their preserved orientations. If the tuffs were water-lain, they are not necessarily associated with a gravity-driven flow, and accordingly the fronds were interpreted as having been felled by basin contour-parallel currents (Wood et al. 2003). However, if the smothering ashes are a product of turbidity flows, then it is likely that the fronds were felled by these same flows (Matthews et al. 2021). In the specific case of the E surface, however, there is no contention that there is a gravity-driven flow origin for the alignment of fronds on the E Surface (F12 of Wood et al. 2003; Matthews et al. 2021).

### Gravity flows and their expression in the rock record

Turbidites are the lithological record of deposition via sediment-laden turbidity currents, and exist on a continuum with other gravity-driven flows and their deposits (Haughton et al. 2009; Talling et al. 2012). The lack of evidence of fluvial input, together with the slump horizons that occur throughout the Mistaken Point Formation (Wood et al. 2003), suggests that the source of the flows in the Mistaken Point Formation are more likely to be those dominantly sourced from slope failure (slumping), rather than rivers. Therefore, here we focus only on the former. Gravity flow behaviour, and thus classification, is principally driven by two factors: the fraction of cohesive components within the sediment, and the overall concentration of sediments within the flow (e.g. Haughton et al. 2009). Higher sediment concentrations, and higher fractions of cohesive components (clay minerals and reworked muds), act to dampen turbulence at the sediment-water interfaces (Cantero et al. 2012; Talling et al. 2012), and within dilute turbidity currents (Baas and Best 2002; Baas et al. 2009).

The coarse tuff immediately above the E surface could be indicative of particle sorting and winnowing within the more turbulent head of the turbidity current (Sparks and Wilson 1983), while the structure within the rest of the bed is consistent with the hybrid flow model of Haughton et al. 2009 (Matthews et al. 2021). On a broad scale, this mixed/hybrid flow interpretation may be reflected in the increased turbidite thickness within the Mistaken Point Formation (previously interpreted as turbidite ponding, (Ichaso et al. 2007)). Surface weathering, synsedimentary and early diagenetic alteration of the volcaniclastic source for the Mistaken Point turbidites would have produced a clay-rich source sediment (cf. Kiipli et al. 2007), enhanced by addition of deposition from nepheloid plumes or as hemipelagic fallout (cf. Kenchington et al. 2018). This high clay content and high sediment load would have increased the cohesion within the flow and so dampened its turbulence, potentially generating conditions conducive to internal laminar flow, while dilution of the turbidite head likely brought concentrations below the threshold for a laminar-dominated regime (similar to the high-density turbidity or lower density mixed flows of Haughton et al., 2009), with turbulent conditions within the turbidite head.

As a flow moves down a slope, it can change character and concentration, reflected in different depositional products (Houghton et al. 2009). For example, after slumping, entrainment of water rapidly dilutes the head of the turbidite, inhibiting sediment-induced turbulence dampening (Hallworth et al. 1993; Cartigny et al. 2013). In contrast, entrainment of clay-rich material would have the opposite effect. Differential dilution-driven turbulence often manifests as Kelvin-Helmholtz instabilities (Liu and Jiang 2014), wherein turbulent eddies rotate about a horizontal axis orthogonal to the direction of turbidite propagation (see “roll waves” of Cartigny et al. 2013) – important when we are thinking about the processes controlling frond orientation.

## MATERIALS AND METHODS

### Data processing

In this study we used mapped data from the E surface given by Mitchell et al. 2019, supplemented with *Fractofusus* data from Clapham et al. 2003. Mitchell et al. LiDAR scanned the E surface using a Faro Focus 330X to ensure spatial accuracy was maintained over large areas. The LiDAR scans resulted in a 3D surface mesh of 1 mm resolution. In order to get sufficient resolution to resolve taxonomic identity, Mitchell et al also laser scanned the E surface using a Faro Scan Arm v6LLP, resulting in surface meshes of ∼0.050 mm resolution. The high-resolution scanning was done in grids of ∼1m x 1m. Due to large file sizes, these high-resolution scans could not all be viewed simultaneously, so control points were marked in each high-resolution scan, and in the LiDAR scan, enabling accurate combination of the high-resolution scans with the LiDAR surface data (performed using Geomagic 2015). A photomap was created by photographing the specimens along a horizontal and vertical grid, then using Agisoft Photoscan software v1.3.5 to create a photogrammetric render of the surface. The LiDAR scan was then imported into Photoscan, and the photographs aligned on the LiDAR scan to ensure large-scale accuracy. An orthomosaic of the surface was produced within Agisoft PhotoScan, from which the data was collected. The combination of LiDAR, LLP and photogrammetry enabled accurate retention of angle data between photographs, with minimal perspective projection distortion (Mitchell et al. 2019). Specimens were binned into 7 morphogroups: *Beothukis* – a unifoliate, spatulate-fronded rangeomorph, with a short – or absent – stem and holdfast (Brasier and Antcliffe 2009; Hawco et al. 2020); *Bradgatia* – a multifoliate rangeomorph consisting of up to eight primary branches from a central branching point on an inferred holdfast (Boynton and Ford 1995; Flude and Narbonne 2008); *Charniodiscus procerus* – a unifoliate arboreomorph possessing a circular holdfast, elongate stem, and a lanceolate frond (often laterally displaced) without fractal, rangeomorph-style branching (Laflamme et al. 2004); *Charniodiscus spinosus* – a unifoliate arboreomorph with a large ovate frond, lacking rangeomorph-style branching, tipped with an elongate spine, connected to a large holdfast via a short cylindrical stem (Laflamme et al. 2004); *Plumeropriscum* – a multifoliate rangeomorph composed of at least nine primary branches furcating from an elongate cylindrical stem, attached to the substrate by a discoidal holdfast (Mason and Narbonne 2016); *Primocandelabrum* – a multifoliate rangeomorph consisting of a large holdfast, elongate stem, and substantial crown composed of three first order branches (Hofmann et al. 2008; Kenchington and Wilby 2017) and *Thectardis* – an erect conical taxon lacking evidence of a holdfast (Clapham et al. 2004). We used the size and orientation data from Clapham et al. 2003 for *Fractofusus*, a spindle-shaped rangeomorph (Gehling and Narbonne 2007). We identified 18 *Beothukis*, 52 *Bradgatia*, 61 *C. procerus*, 31 *C. spinosus*, 20 *Plumeropriscum*, 47 *Primocandelabrum* and 27 *Thectardis* across 85.42m^2^ of the E surface bedding plane. We supplemented this data with 1497 *Fractofusus* orientation data from Clapham et al. 2003.

### Retrodeformation

The E surface has undergone tectonic deformation so prior to any analyses, retrodeformation needs to be performed to re-engineer the organisms back to their in-death dimensions (Wood et al. 2003). To perform the retrodeformation, we collected the dimensions and orientations of 24 representative, large, discs across the E Surface (Supp. Fig. 1). Utilising a constant area retrodeformation method, the principle axis lengths for each disc were extracted. Following the methodology of Mitchell et al. (2015), a regression was fitted to determine the retrodeformation ratio (1.75, which within the confidence interval (1.71±0.08) of Mitchell et al. (2015)), which was applied across the entire E surface. To apply this retrodeformation, the annotated photosquares were aligned and stitched together in Inkscape v0.92.4, and rotated to align the principal axes of the mean disc with the vertical and horizontal axes of the document – thus aligning the eigenvectors of retrodeformation with the axes of the document. From here, constant area retrodeformation can be characterised as a deformation, which can be achieved with shortening and elongation of the vertical and horizontal axes. The retrodeformed surface was then rotated to the original orientation. Overall, the photosquares were shortened by 26.7% along the eigenvector oriented 78.5°, and elongated 36.8% along the orthogonal eigenvector of 168.5° (Supp. Fig. 1). We note that, whilst retrodeformation techniques have the potential to introduce error (Liu et al. 2011), the strong correlation of the regression (R^2^ = 0.86) (Supp. Fig. 1) suggests that our retrodeformation technique is suitable for the spatial scale of the mapped E Surface. The orientation measurements are different for *Fractofusus* because unlike the frondose organisms there is no differentiation between the top and bottom half of the organism, such as a disc. As such, the angles are limited to a 180 ° range of 150 ° to 330 ° with the angle of e.g. 200 ° being equivalent to 20 °.

### Statistical analyses

For each taxon population we performed four tests in R v4.0.4. To test for non-uniform distributions of orientation data we used the Rao’s Spacing Test of Uniformity using the package CircStats v0.2-6 (Agostinelli and Agostinelli 2018), with a *p*-value < 0.05 indicating a non-uniform distribution (Rao 1976). For our data, a significant *p*-value indicates non-random felling of organisms, with some orientations exhibiting a greater abundance than would otherwise be expected from random felling. In order to test for multimodal distributions within angular data we used the Hermans-Rasson test (HR test of Landler et al. 2019) using the package CircMLE v3.0.0 (Fitak and Johnsen 2017). Where multi-modal distributions were found, the mean values for each peak were identified utilising the gaussian finite mixture model-based clustering algorithms of mclust v5.4.7 (Fraley and Raftery 2017). To account for the circular nature of angular data, the density distribution was inspected and split at a minimum to ensure any peaks coincident with 0 ° were not bisected. This split produced a continuous 360 ° density distribution with no assumed peak bisection. When more than one distribution was present (i.e. bidirectional distributions), the data were partitioned into two peaks, whilst unimodal distributions (including those found to be composed of multiple coincident distributions) were left unpartitioned. The circular equivalent to a normal distribution is the von Mises distribution, tested using a Watson’s goodness of fit (Agostinelli and Agostinelli 2018). A statistically significant *p*-value output corroborates a von Mises distribution – where a significant von Mises distribution was found, the models of Schnute and Groot (1992) were employed to test for a variety of modelled orientation scenarios. For bimodally-distributed taxa, the constituent distributions were partitioned and frond lengths cross-compared utilising a Mann-Whitney test. Statistical significance would suggest non-uniform sampling from the same parent population; in essence, the orientation-partitioned data would exhibit different frond length distributions.

In order to investigate the spatial distribution of populations which exhibited significant multi-modal orientations, random labelling analyses (RLA) were used. RLA are a type of spatial point process analysis whereby the position of each point (here fossil specimen) is kept constant, but the label (here the orientation group) is randomly permutated about the points (Illian et al. 2008). As such, RLAs do not directly measure the aggregation or segregation between labels (here orientation patterns), so do not test the processes that resulted in labels, but instead measure the differences in spatial distributions of the labels independently of the positions of the fossil specimens (cf. Mitchell et al. 2018). Spatial distributions are commonly described using pair correlation functions (PCFs) which describe how the density of points (i.e. fossil specimens) changes as a function of distance from the average specimen (e.g. Illian et al. 2008). RLAs assess the differences between two characters (orientation group 1 or group 2) of the populations by calculating variations between PCFs by considering the Difference test and the Quotient test (Wiegand and Moloney 2013). The Difference test is the calculation of the difference the distribution of each group in turn (PCF_11_ is the distribution of group 1 and PCF_22_ the distribution of group 2) i.e. PCF_11_- PCF_22_. These differences test the relative aggregation (or segregation) of the spatial distributions of the orientations compared to each other. If PCF_11_ - PCF_22_ = 1 then the orientation groups are randomly distributed about the surface. The Quotient test calculates how the relative group (Diggle et al. 2005) changed with respect to the total density (i.e. the joined distribution of both group 1 and group 2). The distribution of group 1 relative to the joined groups PCF _1_,_1+2_, and group 2 relative to the joined groups PCF _2_,_2+1_ with the Quotient test as the calculation: PCF _1_,_1+2_ - PCF _21_/PCF _2_,_2+1_. If PCF _12_/PCF _1_,_1+2_ - PCF _21_/PCF _2_,_2+1_ > 0 then group 2 is mainly located in areas with high density of the joint pattern, and group 1 is in low density areas (i.e., group 2 has more neighbours than group 1. If this Quotient is significantly non-zero, then the process underlying the characters is density-dependent. In order to test whether any observed patterns were significantly different from a random distribution we follow Mitchell and Harris 2020 and use two different methods, which are commonly used to establish acceptance or rejection of the null hypotheses for ecological data (e.g. Illian et al. 2008 and references therein): 1) Monte Carlo simulations, and 2) Diggle’s goodness-of-fit test *p*_*d*_, which represents the total squared deviation between the observed pattern and the simulated pattern across the studied distances (Diggle et al. 2005). For each RLA test performed, 999 Monte Carlo simulations were used to generate simulation envelopes around the random PCF difference (e.g. PCF_11_ - PCF_22_ = 0) and the *p*_*d*_ values were calculated using Diggle’s goodness-of-fit test. If the observed test (either Difference or Quotient) fell outside the RLA generated Monte Carlo envelopes and also had *p*_*d*_ < 0.1, then the distributions were found to be significantly different. RLAs were performed in Programita (Wiegand and Moloney 2013).

## RESULTS

For all morphogroups, we found statistically significant non-random distributions using the Rao’s Spacing Test of Uniformity and the improved Hermans-Rasson – (all *p* < 0.01, Table 1). The majority of taxa exhibited a non-von Mises (i.e. non-normal) distributions as per the Watson’s test (Table 1). One *Bradgatia* cohort (*p* < 0.05), and the *Primocandelabrum* stems (*p* < 0.01), and fronds (*p* < 0.05) exhibited von-Mises distributions (Table 1). The von-Mises distributions for *Primocandelabrum* enabled model fitting to the orientation distributions of *Primocandelabrum*, which were found to exhibit bi-modal distributions (Supp. Fig. 2).

**Table 1.** Results of uniformity tests, with 5% significance levels used to indicate rejection of the null models i.e. non-random orientations for the Rao’s spacing and Hermans-Rasson tests and von Mises distribution (normally distributed orientation data) for the Watson’s test.

| Taxa | Measurement | Rao's Test<br>*p*-value | Hermans-Rasson<br>test *p*-value | Watson's Test<br>*p*-value |
| --- | --- | --- | --- | --- |
| *Beothukis* | Fronds | < 0.001 | 0.0001 | > 0.10 |
| *Bradgatia* | Fronds | < 0.001 | 0.0001 | > 0.10 |
|  |  |  |  | < 0.05 |
| *C. procerus* | Stems | < 0.001 | 0.0001 | > 0.10 |
|  | Fronds | < 0.001 | 0.0001 | > 0.10 |
| *C. spinosus* | Stems | < 0.001 | 0.0001 | > 0.10 |
|  | Fronds | < 0.001 | 0.0001 | > 0.10 |
| *Fractofusus* | Length | < 0.001 | 0.0010 | > 0.10 |
| *Plumeropriscum* | Stems | < 0.001 | 0.0001 | > 0.10 |
|  | Fronds | < 0.001 | 0.0001 | > 0.10 |
| *Primocandelabrum* | Stems | < 0.001 | 0.0001 | < 0.01 |
|  | Fronds | < 0.001 | 0.0001 | < 0.05 |
| *Thectardis* | Cones | < 0.010 | 0.0001 | > 0.10 |
|  |  |  |  | > 0.10 |

Analyses of the number of cohorts within each morphogroup orientation distribution varied between 1 and 5 (Table 2, Figure 4). For *C. procerus* and *Plumeropriscum*, the stems and fronds exhibited uni-modal distributions, with similar mean orientations of 195° and 192 ° for *C. procerus* stems and fronds and 178 ° and 177 ° for *Plumeropriscum* (Table 2, Figure 4). The majority (96.77%) of *C. spinosus* stems and fronds exhibited a unimodal distribution (190 ° and 183 ° respectively), with a single outlier orientated at 326 ° for stem and 328 ° for frond (Table 2, Figure 4, Supp. Fig. 3). Similarly, *Primocandelabrum* specimens exhibit a unimodal distribution for their fronds (95.74%, 187 °) and with a minor bimodal component for the stems (89.36%, 183 °; 6.38%, 237 °), with two singleton outliers, and frond at 14 ° and the second with its stem at 119 ° and frond at 98 ° (Table 2, Figure 4, Supp. Fig. 3). *Thectardis* and *Bradgatia* exhibited bi-modal distributions, with different distributions indicated by the mean orientations being notably different between the two groups, in contrast to *Primocandelabrum* and *C. spinosus* (Table 2, Figure 4). The majority of *Bradgatia* specimens (57.69%) formed a cohort with the mean orientation of 15 °, with the remainder (42.31%) within the cohort at 188 ° (Table 2, Figure 4). The majority of *Thectardis* specimens (74.07%) formed a cohort with the mean orientation of 199 °, with the remainder (25.93%) within the cohort at 17 ° (Table 2, Figure 4). The distribution of sampled *Beothukis* specimens formed 5 distinct cohorts (Table 2), with one specimen notably different at 95 ° to the other four cohorts, which had similar mean orientations with the unimodal taxa orientations. The small number of specimens within the *Beothukis* distributions indicates that the relatively high number of cohorts could be an artefact of small sample sizes. *Fractofusus* exhibited a multi-modal distribution, with four cohorts at 172 °, 236 °, 295 ° and 318 ° (Table 2, Figure 4).

**Table 2.** Cohort analyses for each morphogroup orientation distribution. σ indicates the standard deviation of each cohort with respect to the provided mean orientation. Where all cohorts within a population have equal standard deviation (such as *Beothukis*) a single σ is given, and where each σ varies according to the cohort (unequal variance) then a value is given for each cohort. Note that the *Fractofusus* data is taken from Clapham et al. 2003. Note for *C. spinosus* and *Primocandelabrum* the cohorts of one represents outliers, which were not well resolved by cohort-analyses (Supp. Fig. 3).

| Taxa | Measured | N | Mean Orientation (°) | $\sigma$ | Proportion |
| --- | --- | --- | --- | --- | --- |
| *Beothukis* | Fronds | 1 | 95 | 2.45 | 5.56% |
|  |  | 4 | 135 |  | 22.22% |
|  |  | 5 | 161 |  | 27.78% |
|  |  | 3 | 172 |  | 16.67% |
|  |  | 5 | 185 |  | 27.78% |
| *Bradgatia* | Fronds | 30 | 15 | 26.79 | 57.69% |
|  |  | 22 | 188 |  | 42.31% |
| *C. procerus* | Stems | 61 | 195 | 18.89 | 100.00% |
|  | Fronds | 61 | 192 | 30.23 | 100.00% |
| *C. spinosus* | Stems | 30 | 190 | 30.87 | 96.77% |
|  |  | 1 | 326 | NA | 3.23% |
|  | Fronds | 30 | 183 | 24.83 | 96.77% |
|  |  | 1 | 328 | NA | 3.23% |
| *Fractofusus* | Fronds | 312 | 172 | 13.19 | 20.84% |
|  |  | 678 | 236 | 26.02 | 45.29% |
|  |  | 306 | 295 | 13.44 | 20.44% |
|  |  | 201 | 318 | 5.89 | 13.43% |
| *Plumeropriscum* | Stems | 20 | 178 | 16.73 | 100.00% |
|  | Fronds | 20 | 177 | 17.26 | 100.00% |
| *Primocandelabrum* | Stems | 42 | 183 | 13.74 | 89.36% |
|  |  | 3 | 237 | 13.74 | 6.38% |
|  |  | 1 | 11 | NA | 2.13% |
|  |  | 1 | 119 | NA | 2.13% |
|  | Fronds | 45 | 187 | 19.05 | 95.74% |
|  |  | 1 | 14 | NA | 2.13% |
|  |  | 1 | 98 | NA | 2.13% |
| *Thectardis* | Cones | 7 | 17 | 33.00 | 25.93% |
|  |  | 20 | 199 |  | 74.07% |

**Figure 1:**
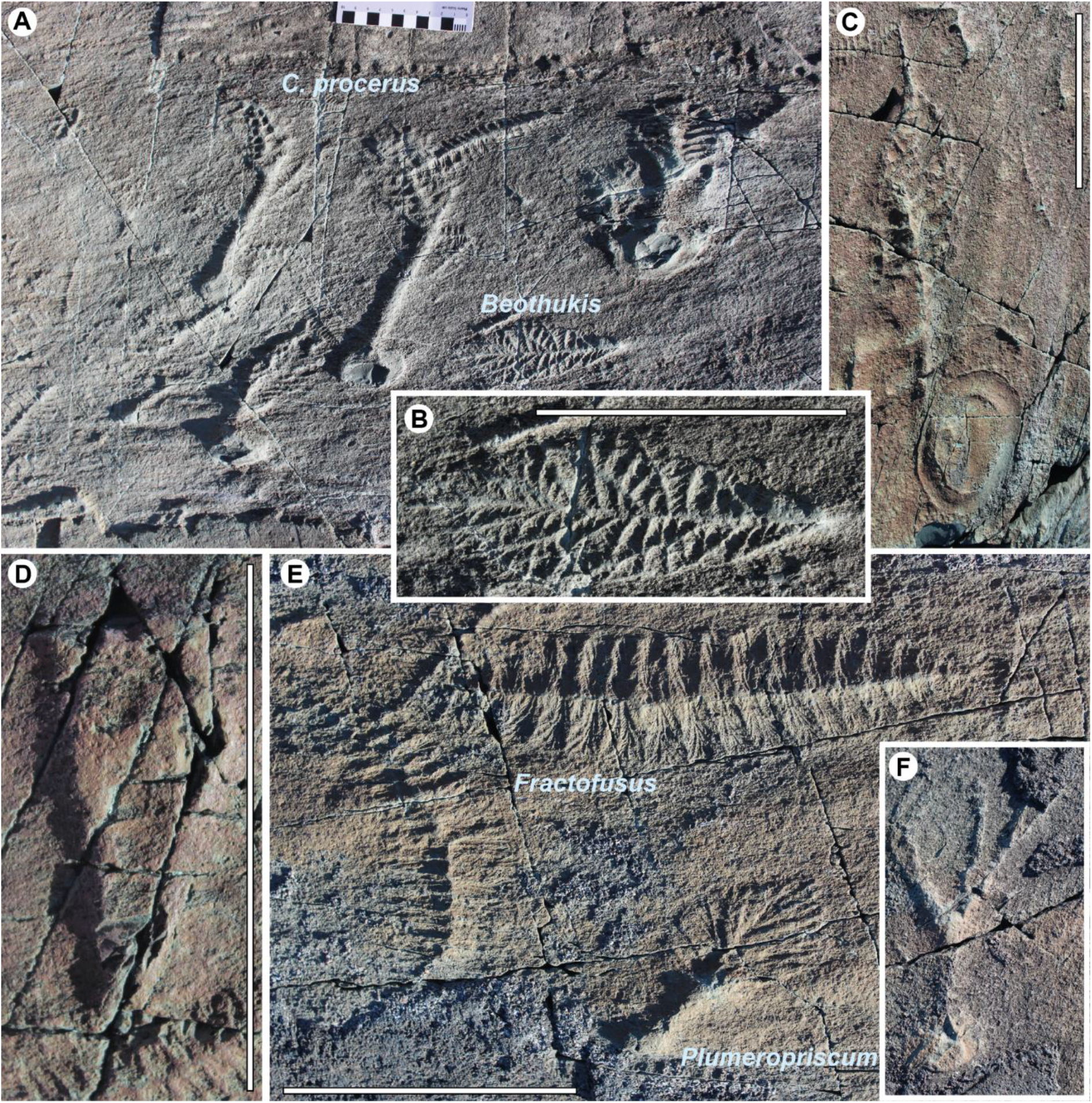
E surface taxa included in this study. A) *Charniodiscus procerus* and *Beothukis* and B) Close up of *Beothukis* C) *Charniodiscus spinosus* D) *Thectardis*, E) *Fractofusus* and *Plumeropriscum* and F) *Primocandelabrum*. Scale bar is 5cm.

**Figure 2:**
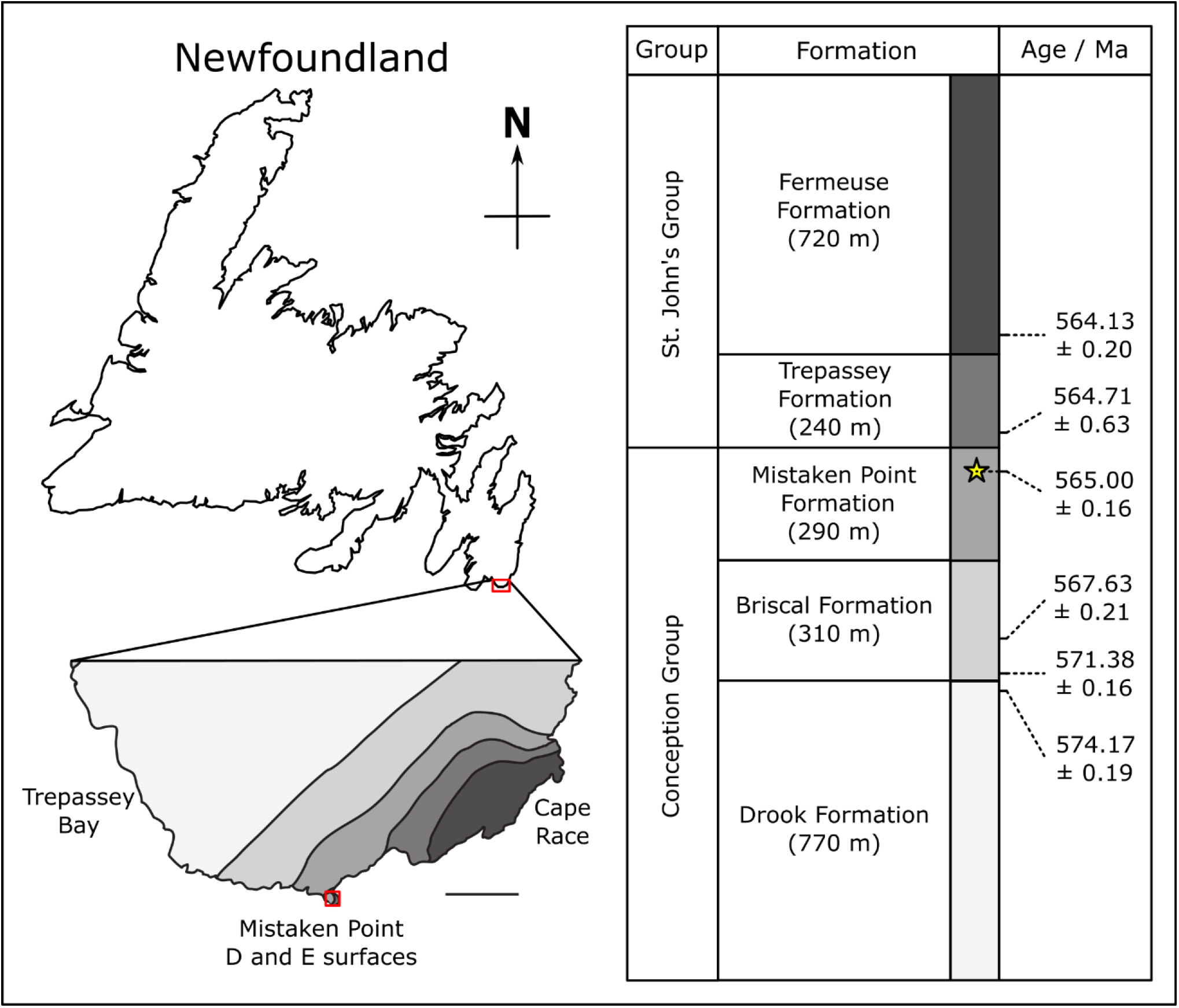
Geological Map after Liu (2016) and Matthews et al. (2021) showing the location of the E surface, Mistaken Point within Newfoundland, Canada, and the stratigraphy and age from Matthews et al. 2021.

**Figure 3:**
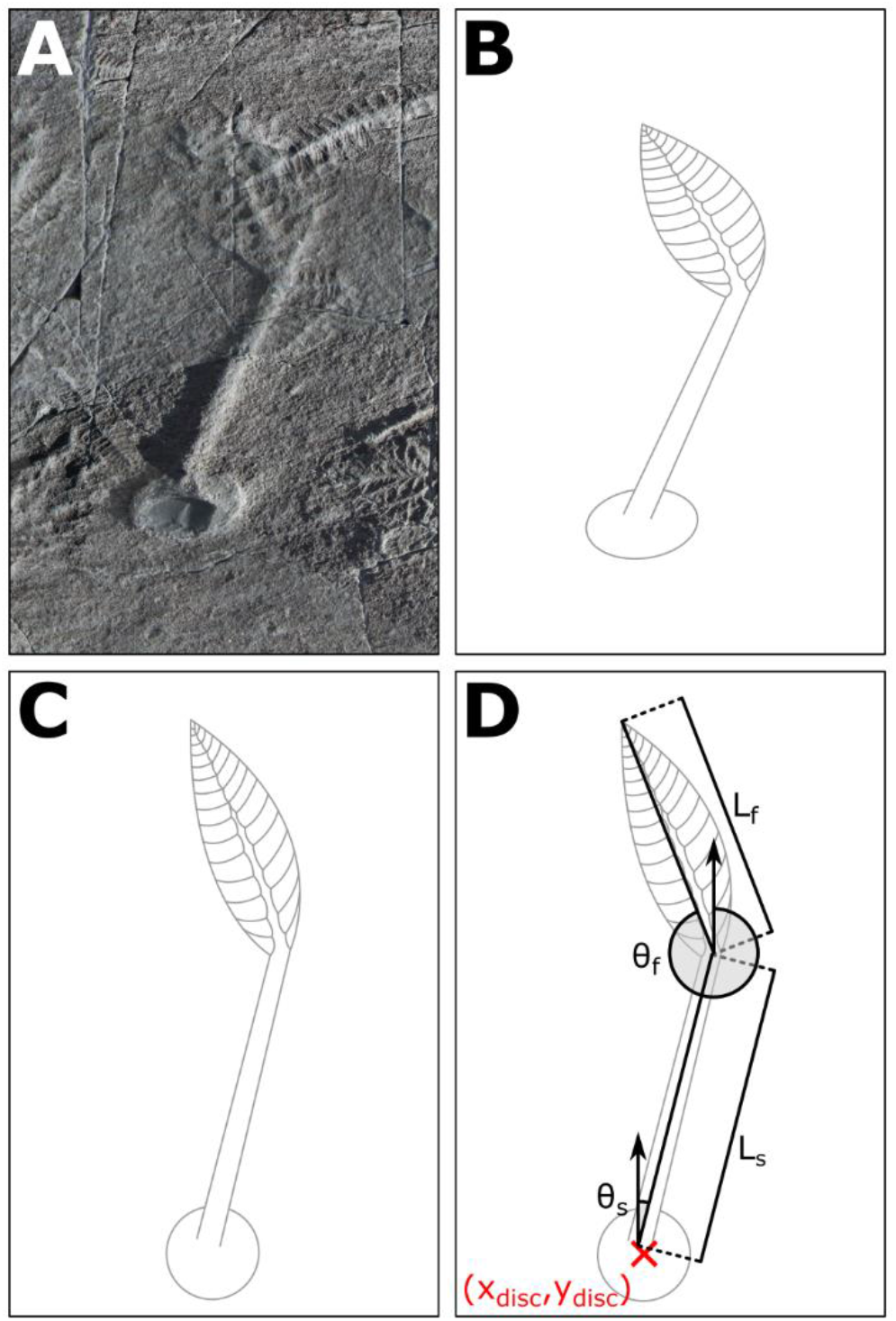
Specimen measurements. An example specimen (A) showing the retrodeformation of the disc B to C, and D) the measurements collected for each specimen. The position of the disc is denoted as (x_disc_,y_disc_); the length of the frond as L_f_; the length of the stem as L_s_; the angle of the frond as θ_f_; and the angle of the stem as θ_s_.

**Figure 4:**
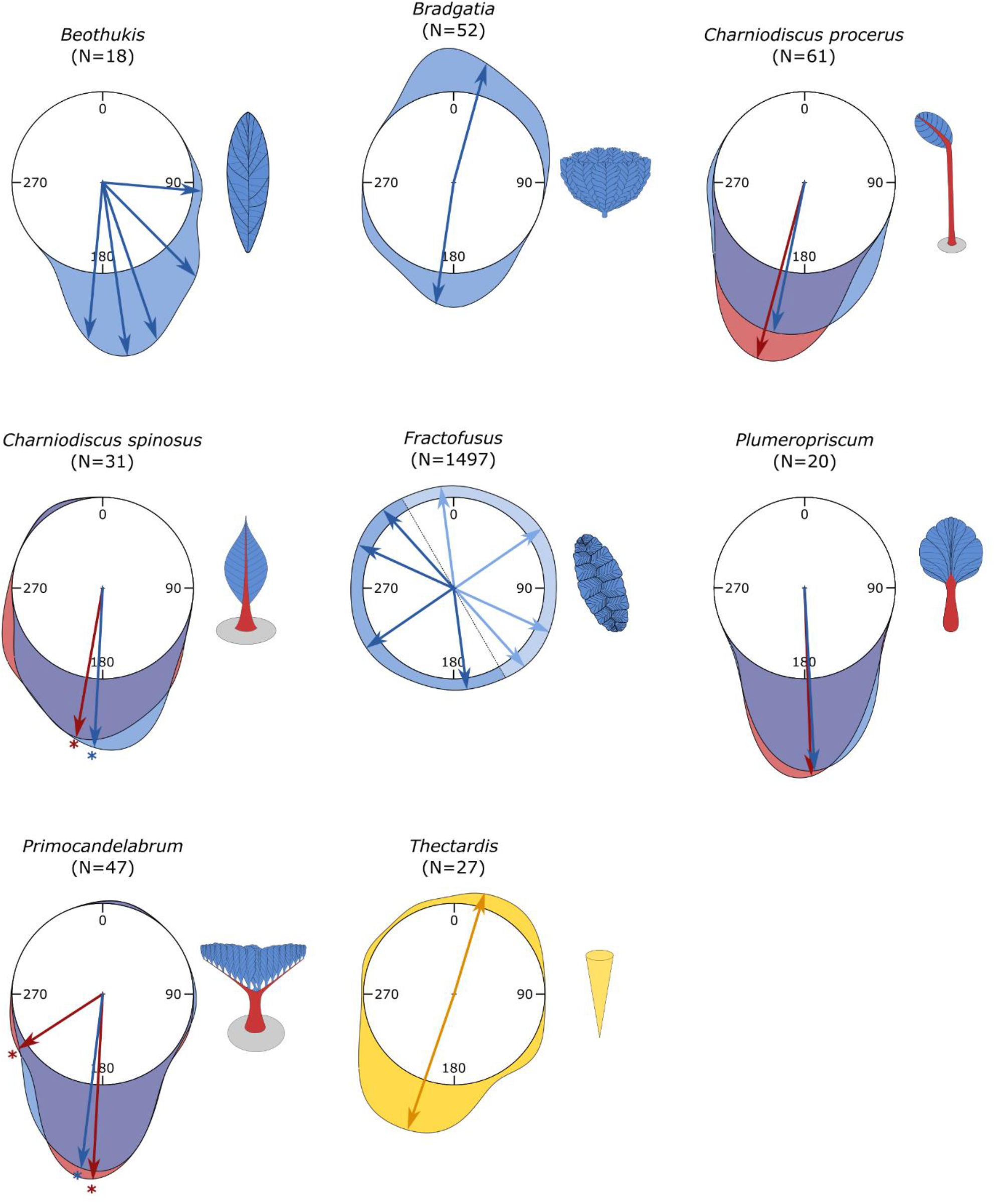
Rose diagrams of the population. Blue indicates frond orientations, Red, the stem orientations and yellow for *Thectardis*. Arrows indicate the mean(s) of the cohort orientation distributions, with starred arrows indicating where outliers have been removed.

Inspection of the distributions in Figure 4 shows that while the number of statistically significant cohorts within each morphogroup varies, the stemmed taxa (*C. spinosus, C. procerus, Plumeropriscum* and *Primocandelabrum*) and *Beothukis* were all orientated in similar directions, while *Bradgatia* and *Thectardis* had a significant proportion of specimens with an antipodal orientation (Table 2, Figure 4). While the mean orientations of the multiple *Beothukis* cohorts were all tightly clustered, showing clear directionality in a single direction, the *Fractofusus* mean orientations were evenly distributed across the range, with no such directionality (Fig. 4).

Analyses of the bimodally distributed taxa found no significant differences mean frond length for *Bradgatia* (*p* = 0.1850), or cone length for *Thectardis* (*p* = 0. 4547) between cohorts. *Bradgatia* was the only taxon that exhibited significant bidirectionality in numbers sufficient for RLA (Fig. 5a). The Quotient test RLA, which describes the relative density dependence of different factors within a spatial population found that there was no density dependence between the two *Bradgatia* orientation groups (*p*_*d*_ = 0.8040, Fig. 5b). The Difference test RLA which tests for the difference between the spatial distributions of the two orientation groups were not significantly different (*p*_*d*_ = 0.4104, Fig 3c.), although the observed difference was close to the outside the simulation envelope which could indicate a larger spatial scale pattern not captured within out data.

**Figure 5:**
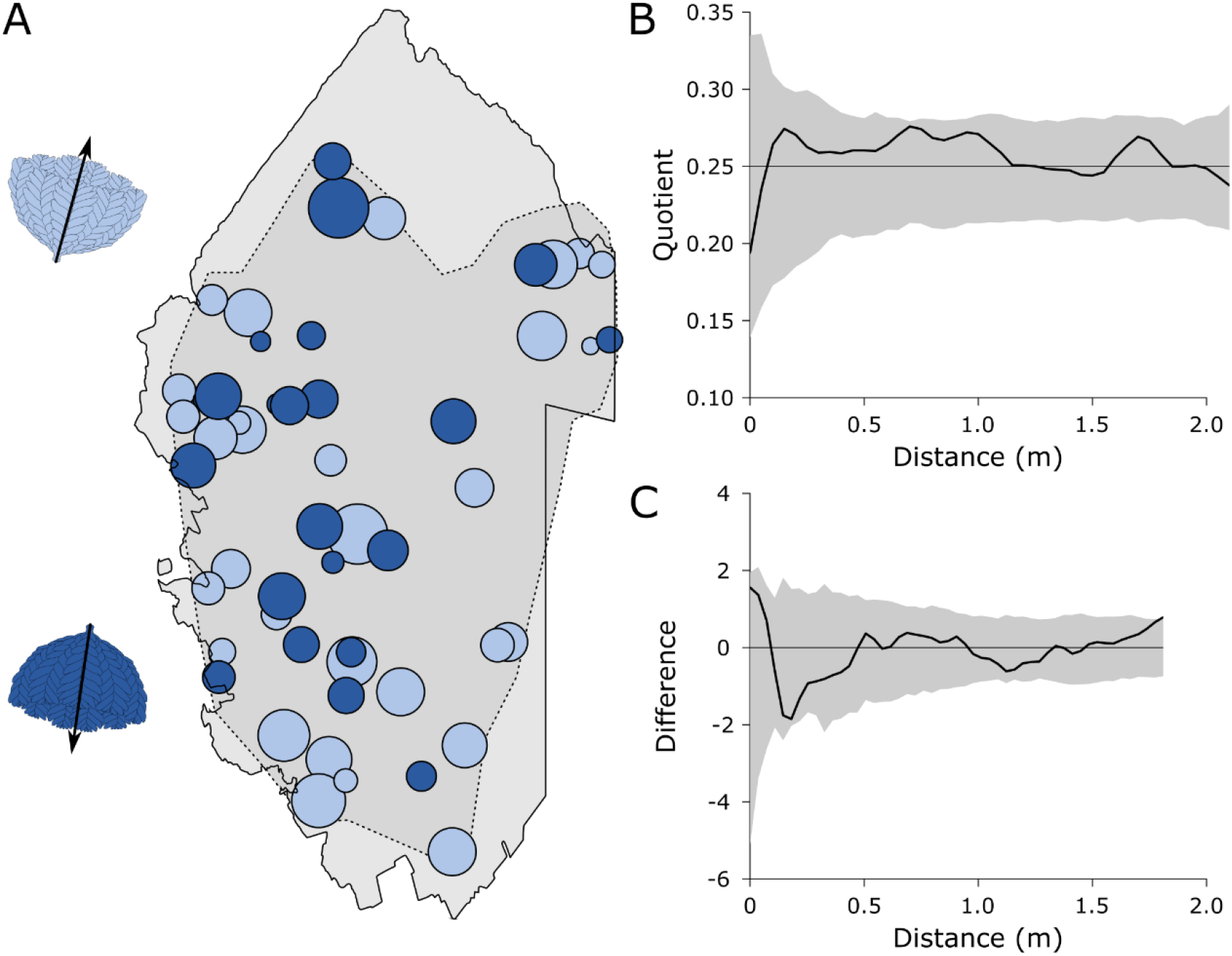
Random labelling analyses for the two cohorts of *Bradgatia*. A) Mapped *Bradgatia* on the E surface, showing the two directions in light and dark blue. The diameter of the circles represents the height of the specimens. Light grey area is the retrodeformed outline of the E surface.

## DISCUSSION

The orientation distributions of fossil specimens are well established as a mechanism to indicate palaeocurrent directions (Toots 1965; Jones and Dennison 1970). The orientation distribution of a given taxon depends on its mode of life, with erect benthic organisms exhibiting strong directionality, in contrast to non-erect organisms which have limited directionality (Toots 1965; Jones and Dennison 1970; Smith 1980; Demko 1995). These explanations of orientation distributions have been used to understand the mode-of-life of Ediacaran taxa, with qualitative examination of 578 *Fractofusus* specimens showing even orientation distribution suggestive of a reclining mode of life (Gehling and Narbonne 2007). In contrast, Ediacaran fronds such as *Charniodiscus* and *Charnia* have been interpreted as erect organisms, due to the morphological similarities to extant benthos such as sea pens (Seilacher 1992; Laflamme and Narbonne 2008; Laflamme et al. 2012) and - crucially - the orientation of these fronds are noted to have a strongly preferred orientation, suggested to be aligned to the contour-parallel current which felled them (Wood et al. 2003; Narbonne 2005; Laflamme et al. 2012). Strongly orientated organisms have been interpreted as erect because an organism attached to the seafloor at a single point will have the majority of its body pulled by the current, orientating it with its long axis parallel to this current. In contrast, if an organism is reclining on the substrate, it will not be subject to such currents, so will not display strong orientations (Gehling and Narbonne 2007; Mitchell et al. 2015).

The relationship between fossil orientation and mode-of-life is pertinent because there has recently been revived debate surrounding the nature of the life habit of the Ediacaran rangeomorphs (e.g. McIlroy et al. 2021). Where fronds have historically been interpreted as displaying a mixture of upright and recumbent lifestyles, recent work has posited that recumbent lifestyles are more likely for some rangeomorph fronds. Orientation analyses allows us to test between these two different life habits in a statistically valid way. This study is the first to quantitatively test the orientation distributions of E surface taxa, and the first to record differences in orientations between the stems and fronds of these taxa. We find significant differences in felling behaviour between the stemless *Bradgatia* and *Thectardis, Fractofusus*, and all other taxa (*Beothukis*, the arboreomorphs, *Primocandelabrum*, and *Plumeropriscum*), which show unidirectional felling all oriented towards the south. *Fractofusus* shows no notable directionality in any direction, whereas orientation distributions of *Bradgatia* and *Thectardis* both exhibit evidence of bidirectional felling. We found no height/sized-based correlations with orientation or outliers. Our results confirm the qualitative results of previous authors (Laflamme et al. 2012, Wood et al. 2003; Narbonne et al. 2005; Gehling and Narbonne 2007) whereby frondose taxa such as *Beothukis, Charniodiscus* and *Primocandelabrum* were erect in the water-column, anchored to the sea-floor, while *Fractofusus* lived close to the substrate in a reclined habit. Our results do not support recent suggestions that the fronds like *Beothukis* reclined on the sediment in life (McIlroy et al. 2021). The orientation distributions we find for *Bradgatia* and *Thectardis* are also consistent with an upright mode of life and felling in a (bidirectional) current. All of our results and interpretations are based on the behaviour of the majority of specimens within a taxon, and confirm the utility of populations of specimens rather than outliers to infer the ecology for the entire population of a given taxon (e.g. Benhadi-Marín 2018). Describing population distributions enables intra-specific variability to be captured, and thus enables comparison between populations. Indeed, it is not possible to compare the orientations of two specimens in a statistically rigorous and robust way without accounting for intra-specific variability, i.e. without quantifying the population behaviour. There are, notably, multiple cohorts within the orientation distributions of *Beothukis*. However, while *C. procerus*, for example, exhibits different mean orientation directionality to *Beothukis*, the 95% confidence interval (as given by two sigma) places all bar one specimen of *Beothukis* (the holotype, oriented at 95 °) the *C. procerus* confidence interval – and indeed, within the 95% confidence intervals of all other southerly-oriented taxa. Thus, the *Beothukis* and *C. procerus* populations do not have significantly different orientations. It is possible of course that the *Beothukis* holotype belongs to a different species than the remainder of the population assigned by us to that taxon based on branching characters. However, recent work on the taxonomy of *Beothukis*, which demonstrates that the holotype is well within all other specimens with comparable morphology, and which were assigned by those authors to that taxon (Hawco et al. 2020) renders this unlikely. Indeed, an outlier of *Primocandelabrum* – whose morphology, and the orientations of the rest of the population, are entirely at odds with a reclined mode of life – is also oriented at 95 °. Our orientation analyses of the *Beothukis* population demonstrates how the holotype orientation is an outlier and not representative of the population. Our results thus confirm an erect lifestyle for *Beothukis* (Wood et al. 2003; Laflamme and Narbonne 2008; Laflamme et al. 2012), contra McIlroy et al. 2020.

*Charniodiscus procerus* specimens – the taxon with the proportionally longest stem of any studied here (Laflamme et al. 2004) – are all oriented south, in a single cohort, with 1 outlier individual oriented antipodally (Supp. Fig. 3). All bar one specimen of *Beothukis* is oriented south, although notably with greater variance than the stemmed arboreomorphs (Fig. 4). All *Plumeropriscum* specimens are oriented south, along with the majority of *Primocandelabrum* (Fig. 4). Two *Primocandelabrum* specimens are oriented in a different orientation, away from the main direction (Fig. 4, Supp. Fig. 3). In contrast, *Bradgatia* specimens are divided almost equally between north and south felling directions (Fig. 4). These data would suggest that there is a correlation between proportional stem length and felling direction, and for the multifoliate taxa, there seems to be a strong correlation between presence of a stem and felling direction. *Thectardis* – with its narrow base and wide top – like *Bradgatia*, also shows a significant portion felled in the northern direction (Fig. 4).

Together, our data suggest that those taxa with bases that are proportionally narrow compared to the widths of their tops (*Thectardis* and *Bradgatia*) show significantly different felling behaviours to those taxa that are more elongate and equal in shape, and that those taxa with the longest and thickest stems show the most consistent felling direction. *Beothukis*, for example, appears to show a sympodial central axis, and has the widest spread of any of the unifoliate and dominantly south-felled taxa. Equally, although *Primocandelabrum* has a sturdy stem, it has a proportionally wide top, and two specimens that are felled at a different angle from the main population. *Charniodiscus spinosus* has a much shorter stem than *C. procerus*, and also has a few specimens that are felled antipodally. The top-heavy morphology of *Primocandelabrum, Bradgatia* and *Thectardis* would presumably induce greater drag compared to the more stream-lined unifoliate fronds, making them more susceptible to felling – and also potentially to adhesion to the matground – with the sturdy stems of *Primocandelabrum* helping to redress this susceptibility in all bar a few individual cases.

Random labelling analyses suggest that these differences are not an artifact of different flow regimes in different areas, indicating that differences in orientations between stemmed and stemless organisms may reflect genuine differences in the effect of flow on stemmed and stemless taxa. Fronds and stems behave differently in flow: at a flow velocity of 0 ms^-1^, both the frond and the stem will be fully upright, with no deflection, but as the flow velocity increases the tubular cross-section of the stem maximises the second moment of area, thus reducing the extent of bending under stress, and so this tubular morphology would serve to reduce the probability of failure via buckling (Wegst and Ashby 2007). Perhaps, because of this morphology the stem impeded felling of stemmed taxa within the enhanced velocities of the turbulent head of a turbidite, according with studies concerning the mechanical properties of stems, for example crinoids and aquatic plants (Baumiller and Ausich 1996; Ming-Chao and Chang-Feng 1996; Luhar and Nepf 2011).

These data support a two-phase model of felling (Fig. 6), corresponding to the different flow regimes within a gravity flow. We infer that during the turbulent head of the flow, most fronds were buffeted by Kelvin-Helmholtz vortices. However, some easily-felled taxa (*Bradgatia* and *Thectardis*) were felled by this turbulence, producing a bimodal distribution of felling orientations (Figs 4, 6). The transition to laminar flow within the body of the turbidity flow led to the felling of most remaining fronds, in a unimodal distribution (Figs 4, 6). In Charnwood Forest, we know that at least some fronds were capable of surviving small-scale disturbance events (Wilby et al. 2015). Wilby et al. focussed on the bimodal population structures of the unifoliate rangeomorph *Charnia*, but documented other, stemmed taxa that were also preserved with a bimodal population structure (*Primocandelabrum, Hylaecullulus* and *Charniodiscus*). Together with our data, this suggests that stemmed and elongate taxa showed greater survivability in high velocity flow. Height in the water column has previously been demonstrated to increase propagule dispersal, and doesn’t appear to provide refuge from resource competition (Mitchell and Kenchington 2018). Our work suggests that stems may have had an additional function – lending greater resilience to felling in turbulent and high velocity flow regimes. These insights hint at potential environmental influences on the morphological composition of Ediacaran communities.

**Figure 6.**
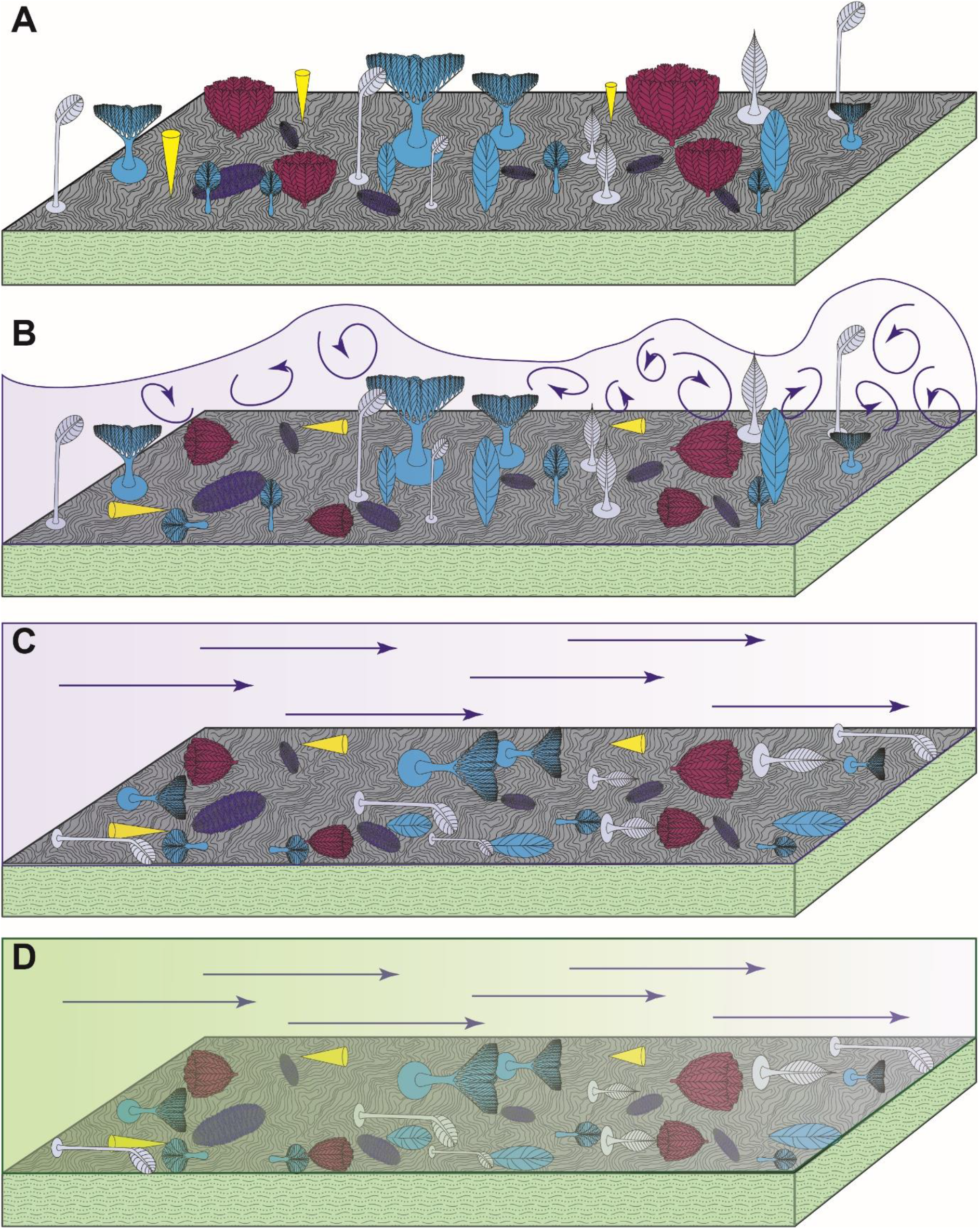
Schematic illustrating the sequence of events that yield the preserved orientation distributions. A) E surface community in life, with all fronds bar *Fractofusus* having an upright mode of life. B) The turbid head of the gravity flow fells organisms with a high centre of gravity, including *Thectardis* and *Bradgatia*, producing a bimodal orientation distribution pattern. C) The laminar tail of the gravity flow fells all other upright fronds on the surface. D) Ash settles out of the flow, and smothers the community with the preserved distribution of orientations. Yellow = *Thectardis*; pink = *Bradgatia*; dark blue = *Fractofusus*; grey = *Charniodiscus procerus* and *Charniodiscus spinosus*; blue = upright fronds felled with a unimodal orientation (*Beothukis, Plumeropriscum* and *Primocandelabrum*).

## CONCLUSIONS

We provide the first quantitative analyses of the orientation of populations of specimens from the Mistaken Point E surface. Our data support traditional palaeobiological models for the life habits of different organisms which lived in this community, with the majority of frondose organisms living upright in the water column while the spindle-shaped *Fractofusus* lived flat on the seafloor. Previous authors have suggested that current type and flow rate may impact community composition, but we demonstrate for the first time how the presence or absence of anatomical features impact survivability in different flow regimes. Specifically, we find that the presence of a stem (and potentially its proportional length) lends greater resilience to turbulent currents. Future work may find that such traits affect the presence and abundance of different morphologies under different environmental conditions, and potentially even the structuring of communities as they experience changing flow conditions.

## Acknowledgements

The Parks and Natural Areas Division (PNAD), Department of Environment and Conservation, Government of Newfoundland and Labrador, provided permits to conduct research within the Mistaken Point Ecological Reserve (MPER) in 2010, 2016 and 2017. Readers are advised that access to MPER is by scientific research permit only. This work has been supported by the Natural Environment Research Council Independent Research Fellowship NE/S014756/1 to EGM and a Natural Environment Research Council grant NE/V010859/1 to FSD. FSD acknowledges support from the Royal Commission for the Exhibition of 1851 and Merton College, Oxford. CGK was supported by Leverhulme Trust (ECF-2018-542) and by the Isaac Newton Trust 18.08(H).

## Author contributions

PBV conceived and designed the project. Analyses were performed PBV and EGM. PBV, EGM and CGK contributed to data collection from photosquares and all authors contributed to the writing up of the final manuscript.

## Supporting information

Supplementary data are available at: (link).

## Data accessibility statement

The original data presented here can be accessed through the Supporting Information.

